# LONG-TERM ECOLOGICAL MONITORING IN INDIA: A SYNTHESIS

**DOI:** 10.1101/2024.08.02.606338

**Authors:** Yadugiri V Tiruvaimozhi, Jimmy Borah, Chandra Prakash Kala, Krushnamegh Kunte, Bharati Patel, K A Sreejith, Rajesh Thadani, Anand M Osuri, Mousumi Ghosh-Harihar

**Affiliations:** Nature Conservation Foundation, Amritha, 1311, 12th Cross, Vijayanagara 1st Stage, Mysuru 570017, India; Krea University, 5655, Central Expressway, Sri City, Andhra Pradesh - 517646, India (;); Aaranyak, 13, Tayeb Ali By Lane, Bishnu Rabha Path, Beltola - 781028, Guwahati, Assam, India (;); Indian Institute of Forest Management, Nehru Nagar, Bhopal, Madhya Pradesh - 462003, India (;); National Centre for Biological Sciences, Tata Institute of Fundamental Research, GKVK Campus, Bellary Road, Bengaluru - 56065, India. (;); ICFRE - Institute of Forest Biodiversity, Dulapally, Kompally SPO, Hyderabad, Telangana - 500100, India (;); Forest Ecology and Biodiversity Conservation Division, Kerala Forest Research Institute, India (;); Centre for Ecology, Development and Research, A-17 Mayfair Gardens, New Delhi - 110016, India. (;); Nature Conservation Foundation, Amritha, 1311, 12th Cross, Vijayanagara 1st Stage, Mysuru - 570017, India (;)

**Keywords:** Questionnaire survey, Permanent plots, Biodiversity, Conservation, Research infrastructure, Tropical ecosystems, LTEM

## Abstract

Long-term ecological monitoring (LTEM) is crucial for understanding ecological processes and responses to environmental change, informing management of natural resources, and biodiversity conservation. Systematic LTEM efforts began in India in the mid-1900s, but there is a lack of comprehensive synthesis of LTEM efforts in the country. Here, we use a wide-ranging questionnaire survey of ecologists coupled with a survey of published literature on LTEM efforts in India to synthesise their thematic and geographical spread, and types of data being collected, and identify key challenges to the establishment and maintenance of LTEM projects in the country. Studies monitor 77 unique subjects across 272 LTEM efforts in India. LTEM efforts are more often located in the Western Ghats and Eastern Himalaya, focused on forest vegetation, and monitoring factors such as abundance, distribution, species richness, and biomass. Regions such as North-Eastern, Eastern, Central, and North-Western India, ecosystems such as grasslands, deserts and wetlands, organisms such as macrofungi, amphibians, and reptiles (other than turtles) are underrepresented. Short turnover times of funding and permits were most frequently reported as hurdles in sustaining LTEM efforts. Data from LTEM efforts have been largely used to produce academic outputs such as journal articles, but have also found use for on-the-ground conservation efforts. Overall, this synthesis can help draw attention to the need for systematic long-term ecological monitoring, help efficient utilisation of existing long-term ecological data, identify regions, species and ecosystem components that are underrepresented in Indian LTEM efforts, foster collaborations, and serve as a starting point to address challenges in sustaining LTEM efforts in India.

## Introduction

Long-term ecological monitoring (LTEM) is any sustained effort to monitor species populations, communities or ecosystem components and processes over decadal or greater timescales. Globally, such efforts have been useful for investigating long-term trends, responses to ecological disturbance, and recovery trajectories of populations, communities and ecosystems, as well as phenology and other biological rhythms (e.g. Bjorkman et al. 2020; Hudson et al. 2022). They have been crucial to assess biodiversity and responses of ecosystem function to disturbances such as habitat degradation, and consequences of, and feedbacks to, climate change and other global changes (e.g. Brown et al. 2001; Bjorkman et al. 2020; Campbell et al. 2022; Hudson et al. 2022; Reed et al. 2022). They have also been critical for evaluating the success of conservation and restoration efforts (e.g. Riginos et al. 2012). LTEM efforts have often helped uncover slow ecological and evolutionary processes, lagged responses, and infrequent events that influence demographic processes, community composition and resilience, and ecosystem structure and function, which short-term ecological studies would have missed (Lindenmayer et al. 2010). Some LTEM studies help tease apart changes over time of interactions between social systems, climate systems and ecosystems (e.g. Jones et al. 2012; Collins 2022). LTEM efforts have also been useful in parametrizing and testing performance of climate and ecosystem models aiming to predict ecosystem responses and global carbon and other dynamics, and in fine-tuning various ecological and conservation methods (Franklin et al. 1990; Alonso et al. 2015; Holmberg et al. 2018). All these have led to a better understanding of long-term ecological dynamics which is crucial for species and ecosystem conservation, and for safeguarding human lives and livelihoods in the face of global change.

LTEM has been a part of ecological research in regions such as the USA and Europe since the mid-1800s and the early 1900s, and have gradually expanded in size and scope (Magurran et al. 2010). For instance, bird and butterfly monitoring efforts in the UK starting from the mid-1900s have expanded into broader population monitoring programmes in Europe and North America. These monitoring programmes subsequently led to the early insights into biological impacts of climate change, including poleward movement of populations and changes in breeding times. They also continue to provide insights into long-term decline of insect and bird populations on the whole and endangered and endemic species in particular (Szabo et al. 2010; Bowler et al. 2019; Pollock et al. 2022). A recent Special Issue in BioScience celebrating 40 years of the Long-Term Ecological Research (LTER) network illustrates the widespread acknowledgement of the utility and importance of LTEM efforts, both among scientists and administrators in the USA (BioScience Special Issue on LTER and Climate Change 2022; Collins 2022; https://lternet.edu). Other efforts such as the National Ecological Observatories Network (NEON) in the USA (https://www.neonscience.org/), the Integrated European Long-Term Ecosystem, critical zone and socio-ecological Research, or eLTER network, and Australia’s Terrestrial Ecosystem Research Network have enabled long-term ecological data collection and tracking ecological changes and their causes at various spatial scales across hundreds of sites (Mollenhauer et al. 2018; https://elter-ri.eu; Guerin et al. 2020; https://www.tern.org.au/about).

There are also several global networks of long-term ecological monitoring sites that leverage large-scale variety in climatic, edaphic and biotic characteristics of ecosystems to address wide-ranging ecological questions. One of the largest is the International Long-Term Ecological Research network (ILTER), which is a network of networks founded in 1993, currently encompassing over 40 member networks, with more than 750 sites across all continents including representation from several tropical countries (https://www.ilter.network; Kim et al. 2018; Mirtl et al. 2018). Intergovernmental efforts such as the Group on Earth Observatories (GEO) have facilitated several national and regional monitoring efforts such as GEO BON to track changes in global biodiversity (https://www.earthobservations.org; https://geobon.org). Others focus on specific ecosystems, such as ForestGEO (https://forestgeo.si.edu; Condit 1995; Davies et al. 2021), Global Ecosystem Monitoring (GEM) network (http://gem.tropicalforests.ox.ac.uk), International Tree Mortality Network (ITMN; https://www.tree-mortality.net) maintained by the International Union for Forest Research Organisations (IUFRO; https://www.iufro.org), and the Global Ocean Observing System (GOOS; https://www.goosocean.org). Recent decades have also seen the birth of several globally distributed experiments, such as Nutrient Network (https://nutnet.org), that have enabled arriving at causal relationships in addition to patterns and trends uncovered by long-term observational studies.

Collective learnings from these large-scale, long-term monitoring efforts have enabled key insights into the challenges and limitations faced by LTEM efforts. These include effectively defining long-term monitoring, developing study designs, iterative adaptation of LTEM efforts to ensure its continued success and utility, balancing trade-offs between the needs of the scientific community and stakeholder expectations, collaboration, partnership and science-to-policy translation (Lindenmayer & Likens 2009; Lindenmayer & Likens 2010; Burns et al. 2014; Kim et al. 2018). Improving the comparability of long-term ecological data and site metadata around the world, reinforcing long-term observation research with long-term experiments to enable making causal inferences, and improving representation of certain ecologies and geographies in global long-term monitoring efforts are also important concerns (Lindenmayer & Likens 2010; Magurran et al. 2010; Mirtl et al. 2018). This is in addition to other common challenges of long-term efforts, including site-level difficulties (e.g. accessibility), data related challenges (e.g. data management), and logistical difficulties (e.g. sustained funding, personnel) (e.g. Burns et al. 2014; Davies et al. 2021).

A major factor limiting the scope and utility of global long-term ecological data is the underrepresentation of tropical and other historically neglected regions, including India, in LTEM efforts and networks (e.g. Mirtl et al. 2018). The relatively small number of LTEM sites and networks in such regions, some from as early as the 1930s (ForestPlots.net), have contributed critical data from some of the most biodiverse regions of the world (e.g. Phillips et al. 1998; Clark 2002; Bauman et al. 2022). While many of these are initiatives by researchers in Europe and North America, some are indigenous to the host countries where native ecological and conservation perspectives are valuable (Pennington & Baker 2021). These LTEM efforts have helped assess long-term changes in ecosystems and ecosystems services, and inform conservation and management strategies, such as the Kenya Long-Term Exclosure Experiment (KLEE, Young et al. 1997; Riginos et al. 2012), Experimental Burn Plots in Kruger National Park (van Wilgen et al. 2007), and the Chamela Region, Mexico (Maass et al. 2005). There are also a few regional and national networks that have helped sustain and expand LTEM efforts in these regions, such the South African Environmental Observation Network (SAEON) supported by their National Research Foundation (https://www.saeon.ac.za).

In India LTEM appears to have emerged from a utilitarian philosophy, focusing on keeping track of forests as sources of trees and animals that were economically or recreationally valuable (Tewari 2016; Jhala et al. 2019). Efforts towards long-term ecological monitoring from an ecological or conservation motivation were probably started in the mid-1900s, and have gathered traction over time (Tewari et al. 2017). Currently in India, there are a few nationally coordinated, government funded and centrally administered long-term monitoring projects such as Project Tiger (https://ntca.gov.in; reviewed in Jhala et al. 2021), and Project Elephant (https://moef.gov.in/project-elephant-pe). These have contributed to country-wide data collection, assessment and conservation implementation. However, there are also several smaller-scale LTEM efforts in India, led by individuals, or groups of ecologists. Most efforts at collating information about such LTEM efforts in India have focused on particular groups of subjects monitored. For instance, Tewari et al. (2014, 2015, 2016, 2017) comprehensively review forest observational studies in India; while several of the efforts outlined in these are still ongoing, many are past efforts with unpublished data (with data also not digitised in many cases). There are also journal articles, reports and other literature on government-led monitoring efforts such as national forest inventory of India (https://fsi.nic.in/about-forest-inventory), long-term research and monitoring of Asiatic lions in Gir (Jhala et al. 2019), and Project Tiger (https://ntca.gov.in). The Indian Long-Term Ecological Observatories (LTEO) Science Plan document and related literature briefly summarise existing, ongoing LTEM efforts in India across ecosystems, but these lists are only indicative of current efforts (https://lteo.iisc.ac.in; Indian Long-Term Ecological Observatories, 2015; Negi et al. 2019). A synthesis of LTEM efforts in India is essential to facilitate better utilisation of existing long-term ecological data, avoid replication of effort, identify regions, species and ecosystem components that are underrepresented, and recognise and address common challenges in sustaining and publishing data from such efforts.

Here, we aim to provide such a comprehensive synthesis of LTEM efforts in India. Specifically, we aim to address the following questions: (a) Where are LTEM efforts in India located? (b) What are they monitoring at various ecological scales? (c) What are the sources of funding for LTEM efforts in India? (d) What are the outputs from LTEM efforts in India? This synthesis also goes one step further to ask ecologists leading and sustaining Indian LTEM efforts about challenges they have faced in maintaining their LTEM projects, as well as in publishing data collected in these efforts. We hope that these first-hand inputs will highlight difficulties faced in long-term ecological monitoring efforts in India and encourage conversations about how to better support and sustain these efforts to serve the goals of species, biodiversity, and ecosystem conservation, as well as the well-being of ecologically vulnerable communities.

## Methods

We considered ‘long-term’ projects as ongoing ones with the intention for long-term monitoring, or, in case of projects that have ended, which were monitored for at least 10 years (Lindenmayer & Likens 2010). Further, we have not considered citizen science efforts, projects carried out by national authorities or institutes (e.g. Project Tiger) and remote-sensing based efforts. This is because the nature of data and challenges associated with national, citizen science or remote-sensing efforts are likely different in character to LTEM studies by individuals or groups of ecologists.

### Questionnaire survey

A survey was sent over email to 113 ecologists involved with one or more long-term monitoring efforts in India. The questionnaire sought responses about the locations of long-term ecological monitoring sites, the subject of monitoring, responses measured at various ecological scales, abiotic and socioeconomic factors that might have also been recorded, sources of funding, outputs from these efforts, and challenges that may have been faced in maintaining the long-term monitoring efforts, and publishing the resulting data (see Appendix I). The list of respondents was compiled by snowball sampling, wherein an initial list of respondents (authors of published LTEM studies) provided information on other long-term projects they were aware of, and so on. This list was expanded by identifying potential respondents based on website searches of about 50 universities and government and non-government organisations involved in ecological research in India, and using a literature survey (see below). We received questionnaire responses of about 69 LTEM projects in India from 46 respondents. The survey was sent to potential respondents after obtaining due clearance from the institutional Research Ethics Committee at Nature Conservation Foundation, Mysuru.

### Literature survey

A literature survey was conducted on 24 March 2022 using the Lens open-access scholarly search platform (https://www.lens.org/). The search terms used were: (“india”) AND (long-term OR observation* OR monitoring OR plot OR decad*) AND (ecosystem OR habitat OR community OR population OR ecolog* OR biodiversity OR behaviour OR dynamics OR change OR conservation OR trend OR rate OR decline OR recovery) NOT (satellite OR GIS OR global* OR agricultur* OR crop OR environment*). We set the ‘Subjects’ list to include Ecology, Evolution, Behavior and Systematics, Nature and Landscape Conservation, Animal Science and Zoology, Aquatic Science, Forestry, Oceanography, Plant Science, Soil Science and Insect Science. This resulted in 4593 publications. This list was supplemented with papers received from our questionnaire responses, and a few from website searches for compiling the respondents’ list. Of these, 54 studies reported data from long-term efforts that had been ongoing for 10 or more years, and these were used for further data extraction. Data extracted from these papers included location of site, duration of data collection, subject of monitoring, and factors measured (in case of 23 projects that were not among the questionnaire responses), as well as information on institutional affiliation of the corresponding authors and sources of funding. Details of a further 13 projects were obtained from literature on institutional websites of ecologists involved in LTEM efforts.

### Data analysis

Data from the questionnaire, literature and website surveys were primarily classified by subject and site of the LTEM efforts. Unique combinations of site and subject were considered one unit, termed as ‘LTEM effort’. ‘Subject’ is defined as the primary focus of the LTEM efforts (e.g. trees in a forest, bird community, or specific species such as the tiger (*Panthera tigris*), or banj oak (*Quercus leucotrichophora*)). Subjects were then categorised into nine ‘groups’ - vegetation, coral, mollusc, fish, insect, reptile, bird, mammal and other. Across subjects, our questionnaire and literature surveys collected data on what is being measured at population, community and ecosystem scales. Given that funding and publications often are for projects that may consist of several sites, the unit at which funding and output related data were summarised and presented is ‘project’, which combines subject- and site- related information. Data were summarised using R version 4.2.1.

## Results

### LTEM efforts in India

We collated 272 long-term ecological monitoring efforts (i.e., unique combinations of sites and subjects monitored) across India, based on data from questionnaire responses, literature survey, and website searches (Fig. 1). Of these, we obtained information on 214 LTEM efforts from questionnaire survey responses, and 37 from the literature survey. Data on the remaining 21 LTEM efforts were obtained from website searches, but are not represented in the questionnaire or literature surveys (Fig. SI1). The majority of studies are based in specific sites (e.g. Rajaji National Park, Agatti in Lakshadweep), while a few are spread over entire regions (e.g. the state of Meghalaya, Spiti Valley, National Chambal Sanctuary). The majority of LTEM efforts are concentrated in a few regions such as the Western Ghats and the Himalaya.

**Fig. 1.**
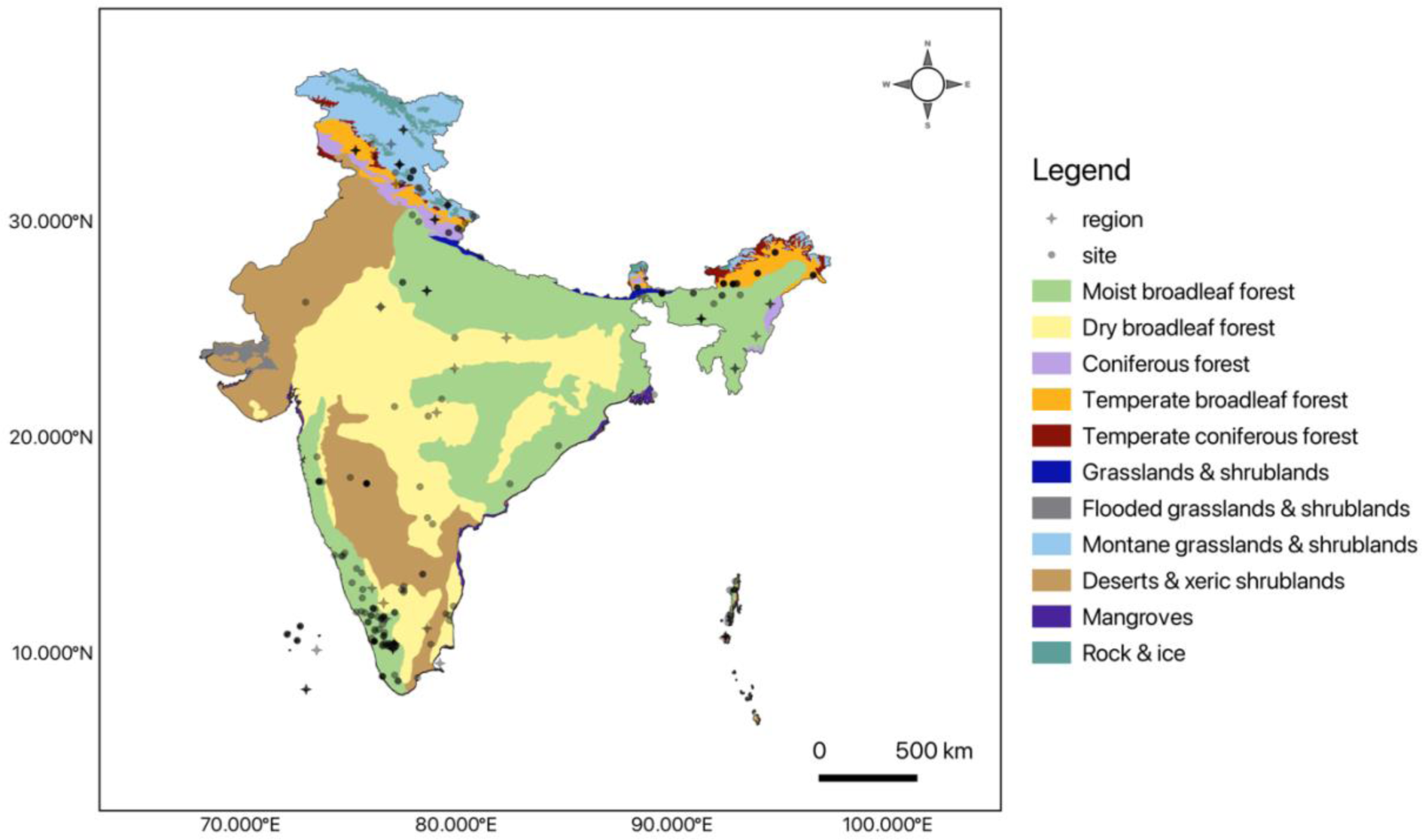
Map of India showing the locations of LTEM efforts. Symbols represent specific sites (circle), or regions (star). Darker symbols are sites / regions with multiple LTEM efforts, and overlapping symbols. Colours on the map are biomes. .

Of 251 LTEM efforts in our questionnaire and literature surveys, 168 have been ongoing for 10 or more years, of which 75 have been monitored for 20 years or more, and 12 for 30 years or more (Fig. 2). However, the frequency of data collection varies widely across cases. While the majority of LTEM efforts have collected data annually, albeit with some gap years where data could not be collected in some cases (i.e., the difference between the age of the LTEM effort and number of years data were collected is 3 years or lesser), some have collected data at lower frequencies (Fig. SI2). For example, in one case data were collected every five years, i.e. four times from a site monitored over 20 years. Overall, 104 efforts have collected data annually for 10 years or more, of which 34 sites have 20 years or more data, and 9 for 30 years or more. Of the 214 LTEM efforts recorded in our questionnaire survey, 200 are still ongoing. All 14 efforts that were discontinued were monitored for 10 or more years, of which 4 were monitored for 20 or more years.

**Fig. 2.**
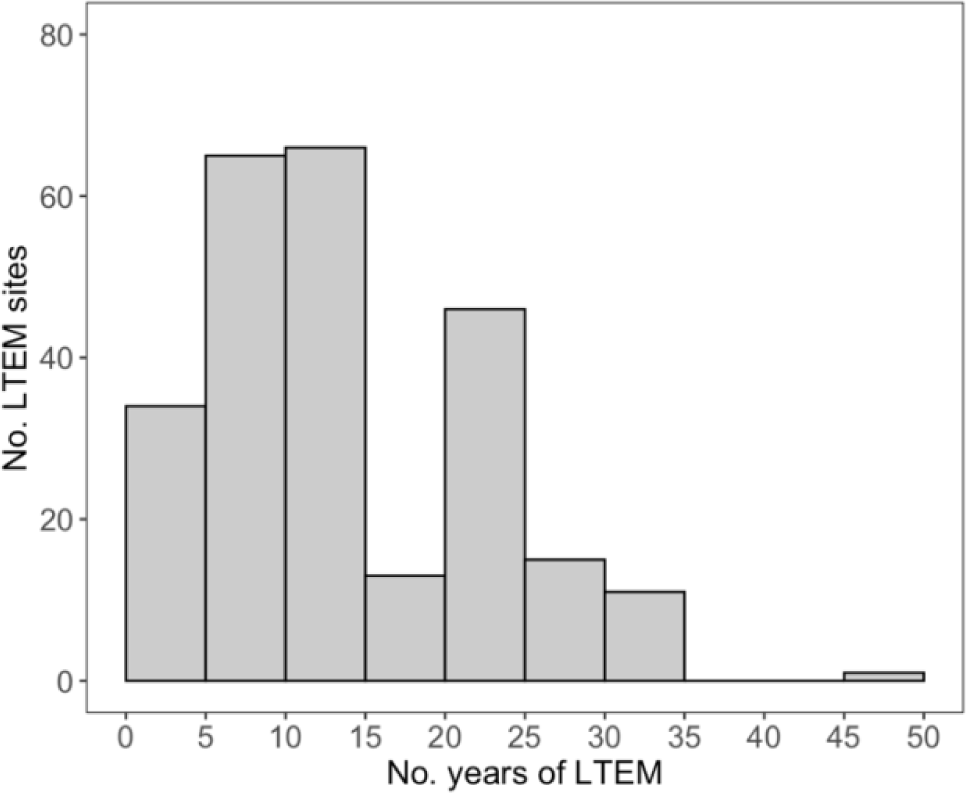
Number of years LTEM sites have been ongoing across India, including all data collection periodicities.

### What are the LTEM efforts monitoring?

There is a vast variety of subjects that are being monitored across sites in India, with 77 unique subjects monitored across 272 LTEM efforts (Fig. 3, Fig SI3). The majority of studies focus on vegetation (97 LTEM efforts, primarily studying forest trees and shrubs). Other commonly monitored groups include birds (53 LTEM efforts, focusing on bird communities in general, or specific groups such as hornbills), mammals (50 LTEM efforts, primarily focusing on tigers, elephants, and several species of primates) and insects (40 LTEM efforts, focusing on butterflies, moths, cicadas and odonates). Fewer studies (i.e. 2-7 LTEM efforts) focus on groups such as coral, molluscs (*Tridacna* species), fish (both marine and freshwater), and reptiles (the majority focusing on turtles), while groups such as amphibians are underrepresented. Even among groups that are well-represented in Indian LTEM efforts, trees and other woody plants are better studied in vegetation LTEM than, for example, marine, wetland or grassland vegetation. Mammal studies tend to focus on tigers, elephants and primates. Insect studies focus on Lepidoptera more than any other group. Macrofungi, many groups of lower plants and invertebrates (including several parasites and key components of marine, soil and other food webs, such as algae, nematodes, annelids, molluscs, arachnids and echinoderms), amphibians, and reptiles other than some groups of turtles do not yet form significant subjects of LTEM efforts in India.

**Fig. 3.**
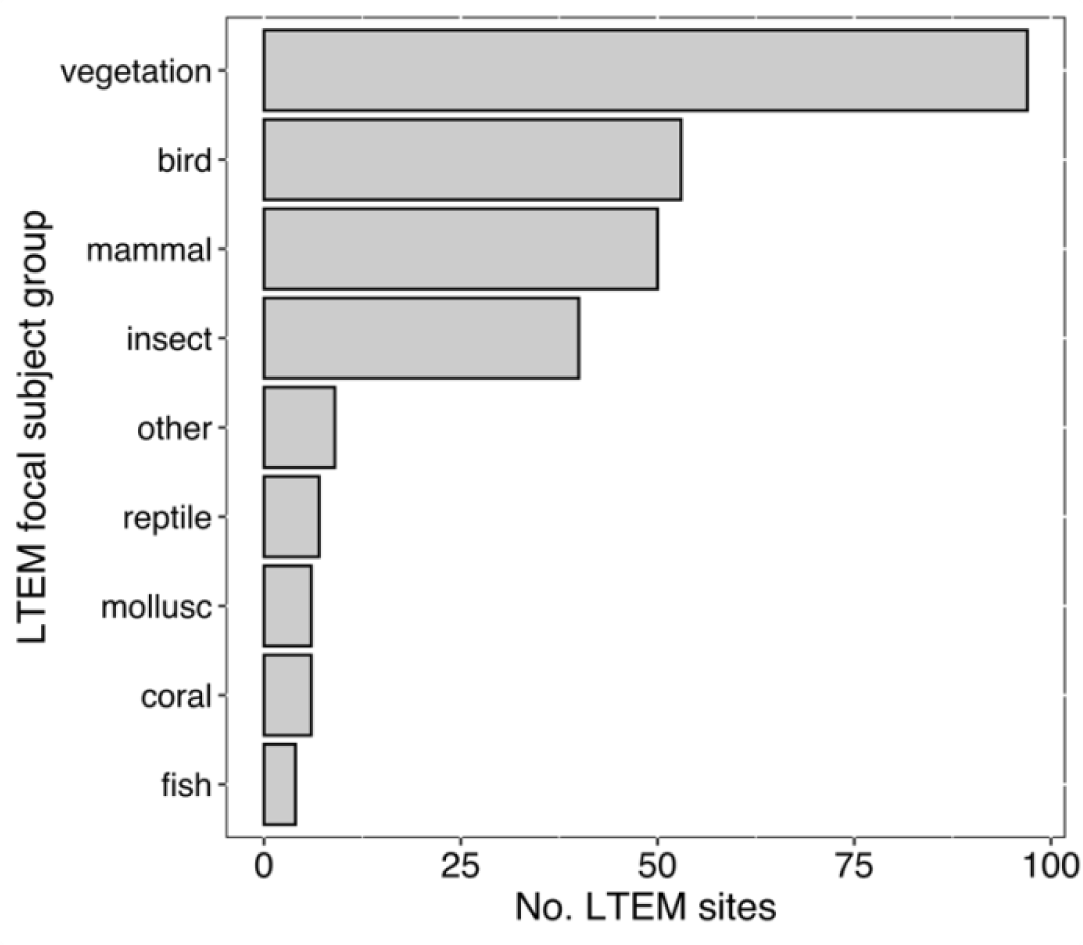
LTEM efforts in India across groups of focal subjects. ‘Other’ includes efforts focusing on biodiversity monitoring in urban areas, ecosystem-scale subjects such as hydrological monitoring, and socio-ecological and socio-economic factors such as human-wildlife conflict and livestock depredation. See Fig. SI3 for details of constituent subjects within each group.

Some LTEM efforts focus on ecosystem-scale subjects such as hydrology, some study urban ecology and biodiversity documenting species across various groups of organisms, and some focus primarily on socio-ecological and socio-economic factors such as human-wildlife conflict and livestock depredation (represented as ‘other’, Fig. 3). Some of the subjects of LTEM efforts are at the community scale (e.g. trees, coral, butterflies), while others focus on specific species (e.g., tiger, rock agama (*Psammophilus dorsalis*)) (Fig. SI3). However, biotic factors that are measured for each of these LTEM subjects may span ecological scales (Fig. 4). Species-level biotic factors are sometimes measured for community-scale subjects; for instance, in a forest monitoring project, phenology data for dominant tree species have been collected. Likewise, community-level factors have been measured for specific species that are LTEM subjects; for instance, a tiger monitoring project may collect data on interactions of tigers with prey species.

**Fig. 4.**
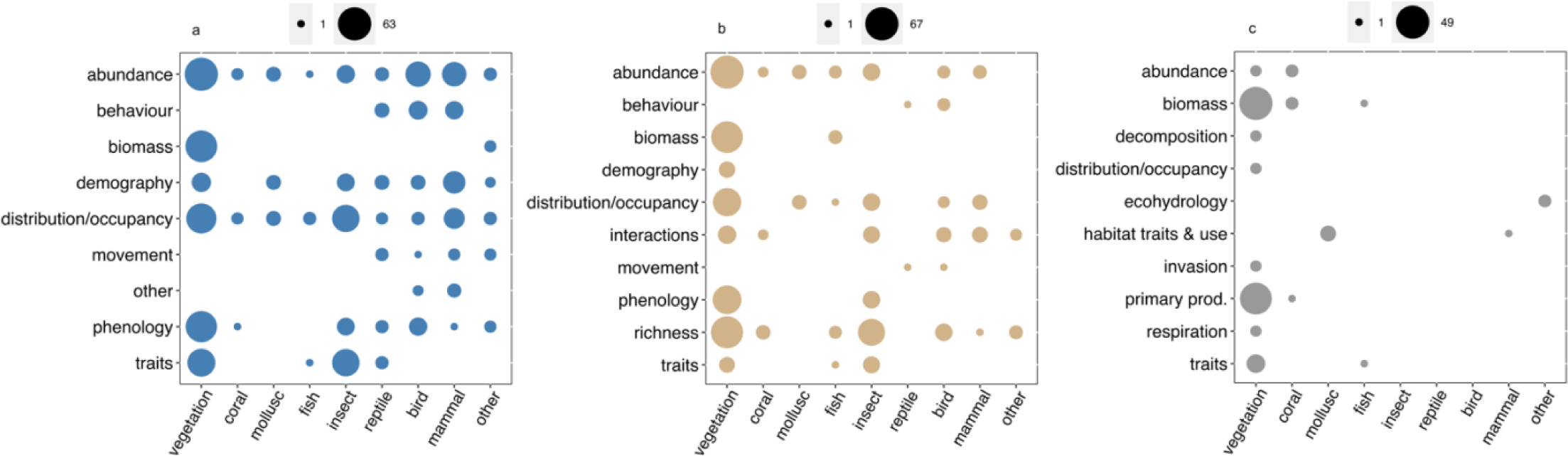
Biotic factors monitored at different ecological scales: (a) species / population scale, (b) community scale, and (c) ecosystem scale at LTEM sites across India. See Table SI1 for details of biotic factors comprising the different groups in the y-axes of the panels.

The distribution of efforts focusing on different subjects reflects the disparity in geographical and ecological coverage of long-term ecological monitoring efforts in the country. Consider, for instance, the three most commonly monitored subjects: vegetation, birds and mammals. Vegetation monitoring is concentrated in the Western Ghats and a few other regions in southern India and in western Himalaya. There is considerably less representation of north-eastern, central and western Indian regions, and consequently of drier regions and ecosystems such as dry forests, grasslands and deserts (Fig. SI4). Bird and mammal monitoring efforts are more evenly distributed across the country. However, some regions in the south such as the Eastern Ghats and coastal regions, as well as the north-western regions of the country are underrepresented (Fig. SI4).

### What are the LTEM efforts measuring?

In our questionnaire and literature surveys, we collected data on biotic factors being monitored at various ecological scales of organisation (Fig. 4, see Table SI1 for details of factors measured within each broad category of factors measured in Fig. 4). At the population / species scale, the most commonly measured factors across all LTEM focal subject groups are abundance and distribution (Fig. 4a). Abundance data (including measures of density, abundance, cover and counts) are collected in 153 of 238 LTEM efforts, and distribution / occupancy data in 129 LTEM efforts. Phenology and traits data are collected in 86 and 83 LTEM efforts, respectively. Phenology data are collected in all groups of LTEM subjects except molluscs and fish. Morphological trait data are collected only for vegetation, fish, insects and reptiles. Demography data (including data on lifespan, mortality causes, population structure and dynamics, and recruitment and survival rates) are collected in 67 LTEM efforts, in all groups except coral and fish. Biomass data are collected in 59 studies, primarily those focusing on vegetation. Behaviour (including data on activity pattern and budget, breeding and reproductive behaviours, habitat preference, and social relationships), and movement data are collected in 31 and 11 efforts, respectively, primarily for birds, mammals and reptiles. A few other kinds of data were also collected at the population / species scale (represented as ‘other’ in Fig. 4a) such as diet preference / composition, and habitat characteristics such as food availability or presence of harmful substances in the vicinity (in an effort monitoring white-rumped vultures (*Gyps bengalensis*)).

Data at the community scale is collected in 173 LTEM efforts, as opposed to 238 efforts that collect data at the population / species scale (Fig. 4b). Species richness is the most commonly collected data at the community scale (127 efforts across all groups except molluscs and reptiles, primarily because mollusc and reptile LTEMs in our survey are species-specific). At least a third of all studies collecting data at the community scale record density / abundance / cover (99 efforts across all groups except reptiles), distribution / occupancy (72 efforts, excluding coral and reptile LTEMs), biomass (64 efforts focusing on vegetation and fish) and phenology data (57 efforts focusing on vegetation and insects). Fewer efforts record data on interactions (such as with prey populations, pollinators, macrofungi, and pests and pathogens; 43 studies), morphological traits (19 efforts on vegetation, fish and insects), demography (9 efforts all focusing on vegetation), breeding and other behaviour, and movement (5 and 2 efforts, respectively, both focusing on reptiles and birds).

As compared to population and community scales, only 68 LTEM efforts record data at the ecosystem scale (Fig. 4c). The majority of these efforts record biomass (53 efforts, primarily focusing on vegetation), and primary production (45 efforts). Fewer efforts record morphological traits (11 efforts focusing on vegetation and fish), abundance (5 efforts focusing on vegetation and coral), and distribution (2 efforts on vegetation). Unlike measurements at the population and community scales, ecosystem scale data are more unique to each project, such as monitoring coral cover as a habitat feature in a study monitoring giant clams. Other factors monitored include ecohydrological response, decomposition, plant respiration, habitat use and plant invasion.

Respondents in our questionnaire survey also reported measuring abiotic and socioeconomic factors alongside the primary focus of long-term monitoring: 65 sites reported monitored abiotic factors, and 49 sites collected data on socioeconomic factors (Fig. SI5). The most commonly monitored abiotic factors are temperature (43 sites) and precipitation (25 sites). Others including various weather parameters, soil quality and texture, pH, water quality and depth and habitat type are monitored in fewer than 20 sites each. Such fine-resolution long-term data on abiotic factors are very valuable especially in the context of understanding ecological responses to climatic and other global changes. Among socioeconomic factors, the most commonly monitored are habitat degradation and fragmentation (19 sites), land use (19 sites), resource use (18 sites), resource extraction (19 sites including fishing and NTFP harvest) and conflict (11 sites). Other factors such as human population, income, livelihood, and cultural practices are monitored in fewer than 10 sites each.

### Funding LTEM in India

We obtained information on funding sources for LTEM in India from 198 projects in our questionnaire and literature surveys. About 90% of these projects have been supported in part by Indian government grants, including schemes such as Long-Term Ecological Observatories (LTEO) and Compensatory Afforestation Fund Management and Planning Authority (CAMPA). International grants (50% of the projects), and state government grants such as by State Forest departments (cited by 43% of the projects), and Universities’ and other organisations’ intramural funding including funding for parts of PhD work (cited by 40% of the projects) are other major sources of funding for LTEM efforts. Corporate Social Responsibility (CSR) and Indian and international philanthropy account for 12% and 9% of funding for LTEM efforts in India, respectively (Fig. 5; see Table SI1 for further details of funding sources). Several respondents in our questionnaire indicated that they did not have any funding for their LTEM effort, or that they used personal funds to sustain (at least part of) their efforts. In many cases, respondents wrote that volunteers, interested students, and local institutions and community members at their field sites contributed time and resources. These play a larger role than is quantifiable for the sustenance of LTEM efforts in India.

**Fig. 5.**
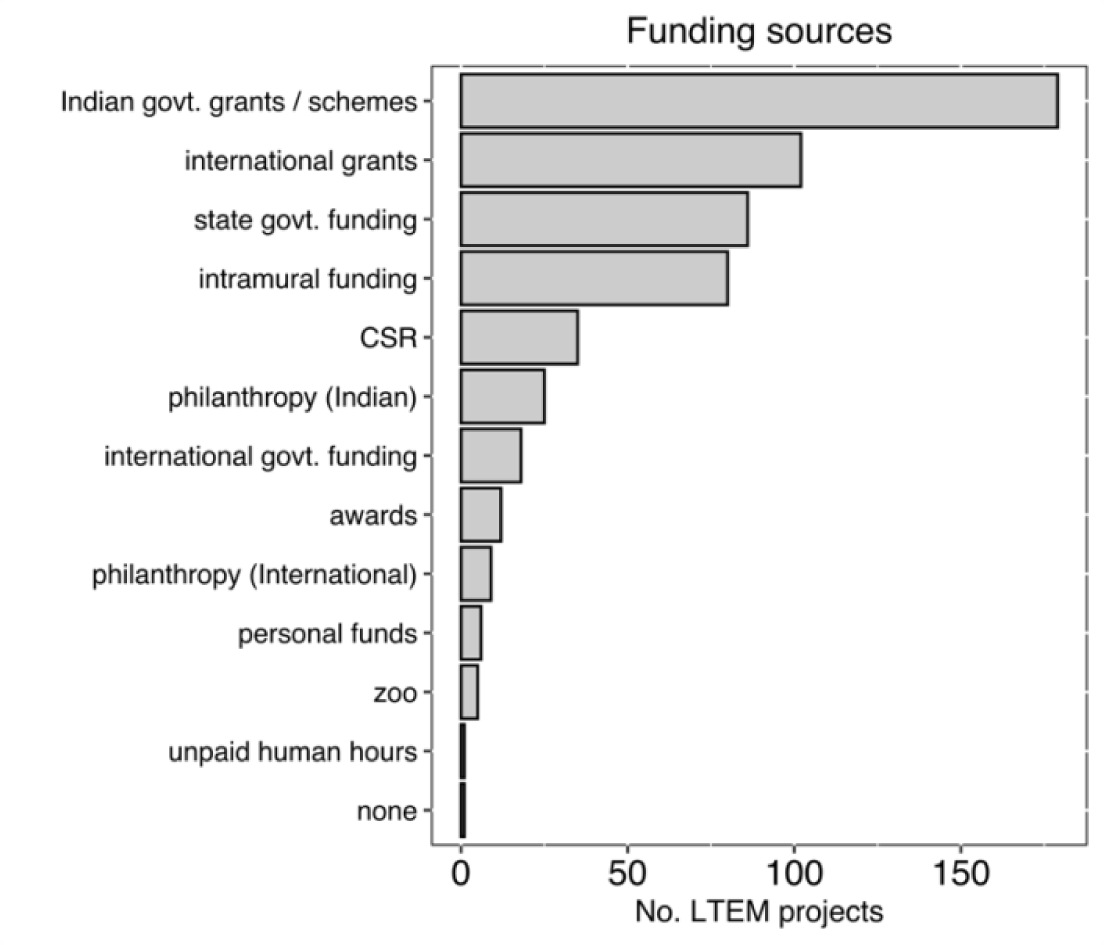
Sources of funding (based on questionnaire and literature surveys).

### Challenges in sustaining LTEM in India

In our questionnaire, we asked respondents about challenges they faced in sustaining their long-term monitoring efforts. We obtained this information for 102 LTEM efforts (i.e., unique combinations of sites and subjects monitored) across India (Fig. 6a). The most commonly cited challenges are that of securing sustained funding (73.5% of LTEM efforts in our responses). As one of our respondents wrote, “The flow of available funds in a timely manner is the single biggest constraint to keep a long-term programme going as the field team has to be supported 24×7 over several decades.” Funding cycles are typically short-term. Many LTEM efforts are sustained by grants raised for other short-term projects in the same sites / region, by linking the monitoring effort to a field-based course in one case, and the monitoring being taken over by the State Forest Department in another. In the words of another respondent, “Grant cycles are short in academia (typically 2-3 years). And it takes a year or two to get a new grant -so one is practically writing for funds all the time!” Funding also links to the other commonly cited problems in maintaining LTEM efforts: the availability of skilled and dedicated human resources for data collection (cited by 54% of LTEM efforts), personnel retention and turnover, and logistics and resources for maintenance of the site and regular monitoring. Personnel turnover is a challenge in multiple ways - new personnel need additional investment of resources for training, and turnover also may impact consistency and continuity of data.

**Fig. 6.**
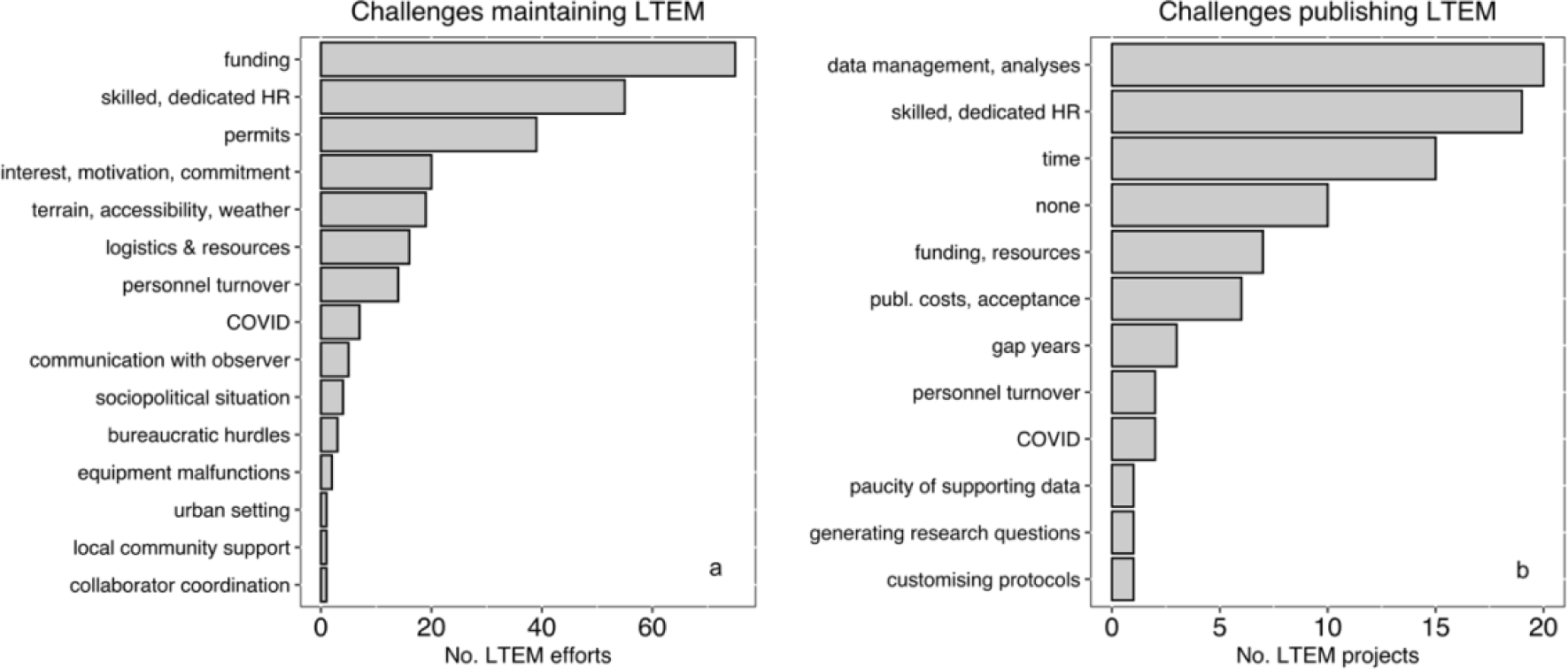
(a) Challenges in maintaining and monitoring LTEM sites, and (b) challenges in publishing data from LTEM efforts (both based on questionnaire responses).

Bureaucratic hurdles are another major challenge in maintaining LTEM efforts. Of these the most cited is the availability of permits from government agencies such as state forest departments for access and data collection in field sites such as protected areas (38% of LTEM efforts), which, like funding, is typically granted only for shorter timescales. This sometimes creates discontinuity in the monitoring effort, and loss of key data. To quote one of our respondents, “One of the biggest hurdles has been the timely acquisition of research permits and the disbursement of funds. This is particularly felt when we are unable to collect regular data or maintain permanent plots which require frequent maintenance and skilled personnel. Additionally, it also affects the monitoring of sudden disturbance events like coral bleaching or disease outbreaks which happen within short time intervals.” Some respondents cited apathy, opposition and red tape at Government / forest department and institutional levels as key hurdles. A case cited by a respondent is where an institute’s general financial rules dictated that the rates for field work assistance are equal irrespective of the level of effort required (for field work at altitudes of 500 m vs. 4000 m, in this case).

Site accessibility, terrain, and weather, as well as socio-political situation in some areas impede long-term ecological data collection (Fig. 6a). LTEM in urban areas have their own unique problems, one of which is the relatively higher human presence in these areas which can encumber long-term data collection. LTEM projects also have unique human resource challenges such as finding people who remain interested and motivated over long periods of time, especially for projects on non-charismatic species. Building lasting relations with local stakeholders, coordinating and collaborating with field-based observers, and, in some cases, with multiple researchers involved with similar monitoring efforts were also cited as concerns.

### Challenges with long-term monitoring data, and products from LTEM efforts in India

We asked our questionnaire respondents about challenges they face in publishing LTEM work, and obtained responses from 47 LTEM projects. Here, too, skilled human resource is a limiting factor (cited by 40.4% of LTEM projects). This links to other difficulties mentioned by our questionnaire respondents, such as the lack of sufficient time (32% of the projects), data management and analyses ability, and paucity of resources and technical expertise for data management and analyses of large, complex, long-term datasets (42.5% of the projects) (Fig. 6b). While about a fifth of our respondents said that they did not face any challenges in publishing LTEM data, some indicated barriers unique to LTEM work. Spatial and temporal scales at which the data are collected is one challenge. One of our respondents wrote, “Publishing is not challenging in itself. Getting such data at relevant scales is the hard part.” Another said, “It is difficult to publish papers on population trends for long lived species with two decadal population data”. Manuscript acceptance is another key hurdle, with journals’ reasons for not publishing ranging from non-standard protocols, to the LTEMs being ‘regional’ studies. As one of our respondents wrote, “Most of the reviewers do not accept our methods as we have tailored them to collect the most relevant data given limited time, unpredictable weather, threats from crocodiles and logistics constraints.” Another respondent said, “Things like one-off observations of unusual behaviours are possible only in long-term studies. However, though these are important for the understanding of the species, they are not easily publishable.”

Despite challenges, there are several products that have resulted from long-term ecological monitoring efforts. We obtained information on products from LTEM efforts from 69 projects from our questionnaire survey. The most commonly reported outputs in these projects were academic in nature, such as journal articles (77% of the projects), theses (36%), book chapters (22%), and review papers. LTEM work is also heavily used to prepare reports, management plans and policies for state forest departments, funders and other agencies (61% of the projects). The variety of outputs from LTEM work, however, is remarkable, and includes evidence in court cases in the Bombay High Court and Supreme Court of India, YouTube videos, popular articles, blog posts, and field guides (Fig. 7).

**Fig. 7.**
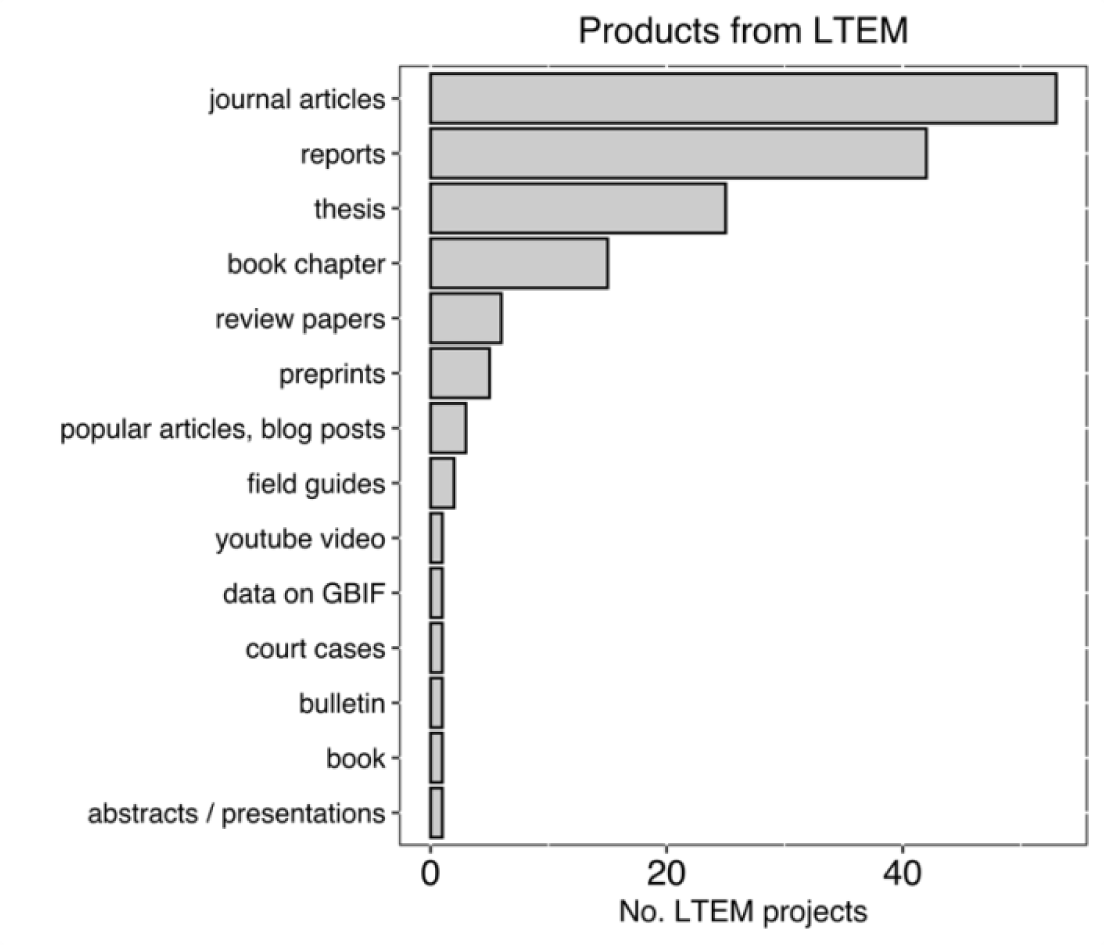
Products from LTEM sites (based on questionnaire responses).

## Discussion

In this synthesis, we used questionnaire and literature surveys to collate data on location, duration, focal subjects, and factors monitored in long-term ecological monitoring efforts across India. LTEM efforts in India, though concentrated in a few regions such as the Western Ghats and the Himalayas, have remarkable variety in the subjects monitored, ranging from seagrasses to trees, and molluscs to mammals. Vegetation monitoring accounts for about 35.6% of all LTEM efforts. Long-term data are collected at multiple ecological scales, with abundance and distribution, species richness, and biomass data being the most commonly collected at the population, community and ecosystem scales, respectively. While grants from the Indian government and international agencies are major sources of funding for LTEM efforts in India, these are often obtained for shorter-term projects in the same landscape given short turnover times of funding cycles and a dearth of dedicated funding for LTEM efforts. Difficulty in securing permits from government agencies is another major challenge in sustaining LTEM efforts in India. Several long term monitoring projects, such as those pertaining to climate change, require an assumption of minimal human disturbance. This makes protected areas a best fit. However, studies in protected areas have access and permit constraints, restricting the spread of subjects monitored in long-term efforts. Despite these challenges, many journal articles and other academic outputs, as well as reports to state forest departments, have resulted from LTEM efforts. These data have also found use in conservation efforts.

A key insight from this synthesis is that while sites for LTEM efforts are distributed across India, some geographies and ecosystems are more well-represented than others. In particular, states in North-Western and Eastern India, and certain ecosystems such as lakes and other fresh-water systems are underrepresented in LTEM efforts in India. Further, monitoring of certain well-studied subjects is regionally concentrated. A probable reason for such geographical biases is the tendency for biodiversity hotspots (e.g. Western Ghats) to attract more attention (Marchese 2015), while logistics and / or accessibility may also play a role. This skewed representation extends to the subjects being monitored as well. This could be due to greater conservation focus and funding available for few charismatic and threatened species (e.g. tigers, elephants). Certain groups are also more conspicuous and easier to identify and monitor (e.g. trees, birds, butterflies), or may be regarded as indicators of ecosystem health due to their association with a diversity of other organisms, or high vulnerability to local microclimate (Bonebrake et al. 2010; Ashe-Jepson et al. 2023). However, other groups of organisms which may be similar ecological indicators (e.g. amphibians, coral, macrofungi) are relatively underrepresented, for reasons including lack of funding, conservation focus and paucity of taxonomic knowledge or difficulty in training in species identification, or their elusive nature requiring more intensive surveys involving multiple methods.

LTEM studies in India have recorded data on several parameters across ecological scales, which have yielded valuable insights into the ecology of a variety of organisms. Long-term abundance, distribution and species richness data, which are the most recorded at population and community scales, have been used to arrive at trends over time and relate them to annual changes in temperature, disturbance and other factors. Examples of such studies include those focusing on population trends of forest trees, molluscs such as *Tridacna maxima*, birds such as the painted stork (*Mycteria leucocephala)* and Indian vultures (*Gyps indicus*), and mammals such as bonnet macaques (*Macaca radiata*) (Erinjery et al. 2017; Apte et al. 2019; Dwevedi et al. 2021; McClure et al. 2021, Singh et al. 2021). Demographic data from long-term monitoring has been used to characterise population recovery for tigers to evaluate conservation effectiveness (Harihar et al. 2020). Changes in community composition such as of coral, trees and birds have been tracked over decadal timescales. In some cases, assessments have been made of their importance in processes such as density dependence driving species coexistence, or community dynamics influencing ecosystem-scale carbon storage (e.g. John et al. 2002; Suresh et al. 2010; Sharma & Singh 2018; Yadav et al. 2018; Marimuthu et al. 2020; Babu et al. 2021). Some studies present longer-term perspectives and results by using chronosequence data, secondary data from the literature, media or forest department and other government records, and surveys and interviews to supplement short- and long-term data from their LTEM sites (e.g. Mallick 2010; Murthy et al. 2022). Some studies have used LTEM data to validate ecological models and predict future trends in population and community trajectories (e.g. Alonso et al. 2015; Antin et al. 2016; Mondal et al. 2016).

Publications from LTEM sites, however, often do not report long-term trends and patterns, but rather results from shorter-term data collected from these sites. While these are perhaps driven by short-term permit and funding cycles, they add to the diversity of results reported from LTEM sites. For instance, several papers discuss taxonomy, compile species lists in the study area, describe distribution / occupancy of the study species, or report discoveries of new species or new behaviours from these sites (e.g. Gross & Price 2000; Kumara & Singh 2004; Apte 2009; Borah et al. 2010; Marathe et al. 2018; Chakrabarti & Jhala 2019; Sharma et al. 2020; Sondhi et al. 2021). Several papers use data and learnings from LTEM efforts to propose improved methods for reliable detection of species and measurement of population densities and abundance (e.g. Karanth et al. 2006; Suryawanshi et al. 2012; Jathanna et al. 2015; Vijayakrishnan et al. 2020). Others evaluate the performance of conservation efforts, or recommend management interventions around the LTEM sites (e.g. Watve 2013; Manchi & Sankaran 2014; Rane & Datta 2015; Bagchi et al. 2019; Raj et al. 2020; Borawake et al. 2021; Hariharan & Raman 2022). Apart from journal articles, products cited in our questionnaire responses also include theses, reports to state forest departments and funding agencies, a YouTube video and evidence in court cases, demonstrating the utility of long-term ecological monitoring efforts in training students and aspiring researchers and conservationists, and as sources of knowledge and evidence enabling conservation efforts.

Some types of data, though commonly collected, are reported more rarely in the literature. For instance, phenology data are commonly collected in LTEM efforts across India, primarily in vegetation studies, but also in efforts focusing on other groups such as birds, insects and coral. However, fewer publications summarise long-term phenology trends or compare long-term patterns in life-cycle timings in plants and other taxa across different sites in India than publications focusing on abundance or richness (Datta & Rane 2013; Ramaswami et al. 2019). Some factors are recorded only by a minority of the LTEM studies in India, such as traits, interactions, behaviour and movement, which may all be key to develop a deeper understanding of various taxa and their long-term dynamics especially in the context of global change (e.g. Sommer et al. 1992; Datta & Rane 2013; Reddy et al. 2019).

In this synthesis, we went one step further to ask ecologists leading LTEM efforts in India about challenges they have faced in sustaining these efforts, with the hope of inspiring conversations to recognise and address these concerns. The survey responses overwhelmingly related either directly or indirectly to the lack of dedicated long-term funding and resources for maintaining the LTEM sites, for data collection, and managing and analysing data. While availability of continuous funding is a challenge in regions such as North America and Europe as well, their importance is recognised leading to better organisation and support systems for LTEMs in these regions (Editorial in *Nature*, 2022). LTEM efforts in India seem to heavily lean on short-term projects in the same area, the dedication of lead ecologists who sometimes spend their personal funds to tide over years with shortage of funds, and the generosity of volunteers, local community members and organisations who contribute their time and resources to sustain these efforts. The existence of LTEM in India is a tribute to the perseverance and dedication of individuals, rather than the result of systemic support by research agencies. Recognition of the importance of broader societal benefits of LTEM, especially to understand how ecosystem change due climate change is impacting ecosystem services and natural resources, may open up newer sources of funding, such as through CSR funding. Sources such as CSR funds may not have been utilised to their potential when it comes to nature conservation in general, and LTEM in particular. Also, the availability of small bridge funding would make it easier to sustain routine monitoring of LTEMs. Once LTEMs are established, maintenance costs are typically lower and all efforts should be made to ensure these initiatives are not abandoned for want of maintenance funds. The Government and research establishment should also prioritise LTEM and it is unfortunate that issues such as lack of permits rank high in challenges in maintaining LTEMs. However, it is to be noted here that this synthesis does not account for other long-term ecological data generated from citizen science efforts (e.g. SoIB 2020, https://www.stateofindiasbirds. in) and dedicated national efforts (such as Project Tiger). These efforts generate data at much larger spatial scales, but face other kinds of challenges. For instance, data quality is a factor that might be of greater concern in citizen-science LTEMs. Efforts such as Project Tiger have faced criticism regarding methodological issues, data availability and access, and political influence on the project (e.g. Karanth 2005; Karanth et al. 2008; Gopalaswamy et al. 2022).

### Key takeaways and future directions

Overall, this synthesis is a first attempt to generate an overview of long-term ecological monitoring efforts in India across focal groups and ecosystems. We hope this synthesis can be a starting point for further collation of information on existing long-term ecological monitoring efforts in India to enable efficient utilisation of existing long-term ecological data, and also to identify regions, species and ecosystem components that are underrepresented in LTEM efforts. We also hope that voices of ecologists leading LTEM efforts as represented here can help initiate conversations to document, recognise and address challenges in sustaining LTEM efforts in India. Some avenues for future conversations and planning for LTEM in India are as follows:

1. *Expanding the scope of LTEMs in India:* Our findings highlight potential areas for future LTEM efforts in currently underrepresented geographies and ecosystems (e.g. deserts, lakes, grasslands), focal groups (e.g. macrofungi, amphibians, bats), and response variables (e.g. traits, interactions, stress responses). There are few LTEM efforts that monitor multiple interacting factors. For instance, as the geographical distribution of bird, mammal and vegetation monitoring in the country shows (Fig. SI4), though plants may affect bird communities and mammal communities, and vice versa, there are few LTEMs that study these interrelationships and interactions. There are also not many LTEM efforts that focus on substrates such as soil or water, processes such as decomposition or CO_2_ efflux, or directly monitor specific global change factors such as invasive species.
2. *Long-term experiments:* Indian LTEM efforts have very few long-term experiments (such as vegetation responses to grazing, nutrient deposition or drought), which are essential to make causal inferences about long-term dynamics inferred from observational studies (e.g. Roy et al. 2023).
3. *Improving ease of long-term monitoring:* Recognition of the importance of long-term ecological monitoring in India should lead to an expansion of sources of funding for such efforts, both via governmental funding sources, and non-government sources such as CSR. Collaboration and support of state forest departments, especially in terms of longer-term permits, are key for the sustenance of such efforts.
4. *Collaboration and knowledge sharing among ecologists:* There is an urgent need for a national collaborative network for long-term ecological monitoring in India. This can facilitate addressing the gaps in current LTEM efforts in India. This can also support data sharing, workshops for training and human resource development, mediating international collaborations, and working towards potential integration with other disciplines for greater impact, such as with climate modellers, and experts in remote sensing and GIS, AI and machine learning.
5. *Applications of LTEM in India:* Conversations and sharing of experiences are necessary about developing long-term collaborations and relationships between researchers from various disciplines, administrators (forest departments), local communities and organisations, and other stakeholders to facilitate efficient utilisation of LTEM data and learnings for the development of informed management and conservation policies, as well as to support, sustain and adapt LTEM efforts for the future.

## Supporting information

Appendix 1: Questionnaire

## Acknowledgements

We are grateful to ecologists leading long-term monitoring efforts in India for sharing their inputs and perspectives via our survey. We thank Abishek Harihar, Aparajita Datta, Kulbhushansingh Suryawanshi, Rohan Arthur, Suhel Quader, T R Shankar Raman and other colleagues at Nature Conservation Foundation for their help in conceiving and developing the idea for this synthesis, collating an initial list of potential respondents, and for their valuable feedback on earlier versions of our questionnaire. We acknowledge funding from Rohini Nilekani Philanthropies and Rainmatter Foundation.

## Appendix

1. Questionnaire
2. Supplementary Figures and Tables

