## Appendix 1: Questionnaire for "LONG-TERM ECOLOGICAL MONITORING IN INDIA: A SYNTHESIS"

This survey is a principal resource for a review of long-term ecological monitoring efforts in India.

The objective of the review is to outline long-term ecological monitoring efforts in India (currently ongoing, or efforts that have at least 10 years' data) - where they are located, what is being monitored, protocols used for monitoring and how they compare with those of other similar monitoring efforts globally, publications (if any) resulting from the research, and sources of funding. We also plan to synthesize findings from some of the monitoring efforts, and identify key aspects to focus on for future long-term ecological monitoring efforts in India. We hope the review will help promote efficient utilization of existing long-term ecological data in India, avoid replication of effort, and identify regions, species and ecosystem components that are under-represented, while also outlining challenges associated with such long-term ecological monitoring in India.

The questionnaire data will supplement data we will gather from literature searches.

We thank you very much for your time and participation.

*Note: By submitting this form you are consenting to your anonymous responses being used for research and conservation projects, and to the data being archived in a public repository.*

### Details of your long-term ecological monitoring effort

**Name of project** *

**Name of respondent(s)** *

**Is this project still ongoing?**

(●) Yes ( ) No

#### Details of your long-term monitoring sites in this project

This information will help us map locations in India with long-term ecological monitoring. This will also help us identify regions that are under-represented in long-term ecology in India.

Please add multiple sites if they are all within the scope of the same long-term monitoring project; i.e. if the same factors are monitored in all of them.

If the sites are very different in what is being monitored, please consider them different 'projects'. We request that you fill separate forms for individual projects.

#### ⊗ Site 1

**Where is your project located?** *

Name of site location, District, State

*Please click on 'Add Site' below to add further sites*

**Year of first data collection**

**Year of latest data collection**

**Number of years data collected** *

+ Add Site

### Details of factors monitored

This section will help us identify the different ecological scales at which long-term monitoring in India has been done, and also the various factors which have been monitored. These data will also help us identify ecosystem components and processes that are under-represented in long-term ecological monitoring efforts in India.

#### If your monitoring is focusing on a few specific species, please provide the name(s) of the species here

Name(s) of species

If your monitoring effort focuses on a single species (such as tiger, or Ficus), or a subset of species in the community (such as a specific set of bird species or plant species in your site), please mention the name(s) of your focal species.

If your monitoring involves more than 10 specific species, please only mention the number of species, and not their names, because their identity would be beyond the scope of our review (e.g.: if 30 tree species in a forest are monitored for phenology, please say '30 common tree species monitored').

#### Population / Species level response factors primarily monitored

- ☐ Distribution / Occupancy
- ☐ Phenology
- ☐ Movement
- ☐ Biomass
- ☐ Demography
- ☐ Interactions
- ☐ Population density / abundance / cover
- ☐ Traits
- ☐ Behaviour
- ☐ Other

Please select all that are relevant at this ecological scale. Please select only factors which are being monitored long-term.

**Community level response factors primarily monitored**

- [ ] Richness
- [ ] Phenology
- [ ] Distribution / Occupancy
- [ ] Biomass
- [ ] Movement
- [ ] Interactions
- [ ] Density / Abundance / Cover
- [ ] Traits
- [ ] Behaviour
- [ ] Other

Please select all that are relevant at this ecological scale. Please select only factors which are being monitored long-term.

**Ecosystem level response factors primarily monitored**

- [ ] Primary production
- [ ] Traits
- [ ] Biomass
- [ ] Decomposition
- [ ] Interactions
- [ ] Other

Please select all that are relevant at this ecological scale. Please select only factors which are being monitored long-term.

**The above lists are not exhaustive. Please use this box to fill in any other factors monitored at the above ecological scales. Please separate individual factors by semi-colons.**

**Abiotic factors monitored**

- [ ] Temperature
- [ ] Water flow
- [ ] Precipitation
- [ ] Soil moisture
- [ ] Wind
- [ ] Soil nutrients
- [ ] Light
- [ ] Soil texture / composition
- [ ] Air quality
- [ ] pH
- [ ] Water quality / nutrient content
- [ ] Fire frequency / intensity
- [ ] Water depth
- [ ] Other

Please select all that are relevant. This is for us to get an idea of abiotic factors that you may also be monitoring in the long-term in your sites.

**Number of years abiotic factors data collected (if this is different from the number of years data were collected for the primary monitoring efforts at each site).**

**Human / socio-economic / other factors monitored**

- [ ] Human population
- [ ] Income / livelihood
- [ ] Conflict
- [ ] Land use
- [ ] Resource use
- [ ] Resource extraction
- [ ] Hunting
- [ ] Habitat fragmentation
- [ ] Cultural practices
- [ ] Other

Please select all that are relevant. This is for us to get an idea of human / socio-economic / other factors that you may also be monitoring in the long-term in your sites.

**Number of years human / socio-economic / other factors data collected (if this is different from the number of years data were collected for the primary monitoring efforts at each site). Untitled**

**If you have anything further to add about any of the factors you have monitored, please leave a note here**

**Publications and other manuscripts these data have been used for**

- [ ] Journal articles
- [ ] Review papers
- [ ] Book chapters
- [ ] Theses
- [ ] Reports (to FD, Grant Agency etc.)
- [ ] Preprints (on bioRxiv or similar)
- [ ] None so far
- [ ] Other

Please select all that are relevant

**Are there any other data that were collected alongside the primary objective of your study, or shorter-term studies that emerged from your long-term monitoring effort? Do you have publications arising from these data?**

For instance, if you have camera trap data for tigers, which was the primary focus of your study, you might also have camera trap data for other animals.

**Details of protocols used for the data collection**

This can include one or more of: a brief description, a link to a website with description, or a reference to protocols of similar monitoring efforts (e.g. RAINFOR), or reference to any publication (not necessarily arising from your work) which has a description of the protocols you are using.

**Sources of funding**

- [ ] Indian government funding (DST, DBT, CSIR etc.)
- [ ] State government funding (e.g. Himachal Pradesh Forest Department)
- [ ] International government funding
- [ ] University / employer intramural funding
- [ ] National grants
- [ ] International grants
- [ ] Corporate Social Responsibility (CSR)
- [ ] Philanthropy (Indian)
- [ ] Philanthropy (international)
- [ ] Awards
- [ ] Other

Please select all relevant sources of funding from the list above. Alternatively, you can also just type in the sources of funding in the textbox below (e.g. DBT, Rufford Grant, CSR). You can also select from the list and leave further details in the textbox below, if that is convenient for you.

**Please add any further details of your sources of funding here**

**If there is anything else you would like to add about this project, or tell us about other long-term monitoring projects you are involved with, please leave a note here**

**What have been the biggest constraints or challenges in keeping your long-term ecological monitoring efforts going?**

This will help us identify challenges associated with conducting long-term ecological monitoring in India. This can include factors such as funding, personnel availability, permits, field sites being difficult to reach, equipment malfunctions, COVID etc. etc.

If you have not faced any constraints or challenges, you can say that too.

**What have been the biggest constraints or challenges in publishing your data from these long-term monitoring efforts?**

This will help us identify challenges associated with publishing data from long-term ecological monitoring efforts in India. This can include factors such as funding, personnel availability for data entry, maintenance and analyses, data loss, gap years, difficulty in getting publications accepted etc. etc.

If you have not faced any constraints or challenges, you can say that too.

**Please add references to publications or other outputs from your long-term monitoring effort (if any) to help us understand your work better**

**You can add pdfs of any of the above references here (up to 5 files)**

Upload or drag files here.

Submit

Please do not submit passwords through Cognito Forms.

Powered by Cognito Forms. Try It Now - cognitoforms.com
