## Appendix 2: Supplementary Tables and Figures for "LONG-TERM ECOLOGICAL MONITORING IN INDIA: A SYNTHESIS"

Yadugiri V Tiruvaimozhi*, Jimmy Borah, Chandra Prakash Kala, Krushnamegh Kunte, Bharati Patel, K A Sreejith, Rajesh Thadani, Anand M Osuri, Mousumi Ghosh-Harihar

**Corresponding author*

Author details:

Yadugiri V Tiruvaimozhi, Nature Conservation Foundation, Amritha, 1311, 12th Cross, Vijayanagara 1st Stage, Mysuru 570017, India. *Current address:* Krea University, 5655, Central Expressway, Sri City, Andhra Pradesh - 517646, India (; ORCID: 0000-0003-1159-277X)

Krushnamegh Kunte, National Centre for Biological Sciences, Tata Institute of Fundamental Research, GKVK Campus, Bellary Road, Bengaluru - 56065, India. (, ORCID: 0000-0002-3860-6118)

**Table SI1** Details of data comprising the biotic factors measured at different ecological scales (see Fig. 4). The questionnaire responses on biotic factors measured were grouped into broader categories for Fig. 4. The details of what these categories include is given below.

| Ecological scale | Biotic factors | Questionnaire responses on biotic factors measured |
|---|---|---|
| Population / species | abundance | density / abundance / cover, counts at roost sites |
| | behaviour | behaviour, activity pattern and budget, breeding, habitat preference, reproductive behaviour, social relationships, survival rate |
| | biomass | biomass |
| | demography | demography, lifespan, mortality causes, population structure and dynamics, recruitment rate |
| | distribution/occupancy | distribution / occupancy, detection / non-detection |
| | movement | movement |
| | phenology | phenology |
| | traits | morphological traits |
| | other | diet, diet composition, diet preference, food availability, prevalence of harmful substances in the vicinity |
| Community | abundance | density / abundance / cover |
| | behaviour | breeding |
| | biomass | biomass |
| | demography | demography |
| | distribution/occupancy | distribution / occupancy |
| | interactions | interactions, interactions with prey populations, macrofungi associations, pollinator abundance, disease and pest incidence |
| | movement | movement |
| | phenology | phenology |
| | richness | richness |
| | traits | morphological traits |
| Ecosystem | abundance | density / abundance / cover |
| | biomass | biomass |
| | decomposition | decomposition |
| | distribution/occupancy | distribution / occupancy |
| | ecohydrology | ecohydrological response |
| | habitat traits & use | coral cover, habitat use |
| | invasion | presence of invasive species, plant invasion |
| | primary production | primary production |
| | respiration | respiration |
| | traits | morphological traits |

**Table SI2** Funding sources acknowledged in 52 publications from LTEM efforts in India, from our questionnaire and literature surveys and website searches (See Methods for details). MoEFCC, India, was cited in 9 publications, Rufford Small Grants Foundation in 6, and DST, India in 4. The rest are cited in 1-3 publications each.

| SI | Funding source | SI | Funding source |
|---|---|---|---|
| | **Indian govt. grants / schemes** | | **Intramural funding** |
| 1 | Central Zoo Authority, Government of India | 38 | DBT-IISc |
| 2 | DBT, India | 39 | Pondicherry University |
| 3 | DST, India | 40 | Ramón y Cajal Fellowship |
| 4 | DST-FIST | 41 | Wildlife Institute of India |
| 5 | DST-SERB | | |
| 6 | ESSO, Ministry of Earth Sciences, India | | **Foundations and philanthropy** |
| 7 | ICAR, India | 42 | Ford Foundation |
| 8 | ISRO-STC | 43 | LEAD International, UK |
| 9 | J.C. Bose Fellowship, SERB, Government of India | 44 | Alan and Patricia Koval Foundation |
| 10 | Ministry of Education, Culture, and Social Welfare, India | 45 | Gordon and Betty Moore Foundation |
| 11 | MoEFCC, India | 46 | Iain Allan, Tropical-Ice Safaris |
| 12 | Man and the Biosphere Programme, MoEFCC | 47 | Liz Claiborne-Art Ortenberg Foundation, New York |
| 13 | N-PDF, DST-SERB | 48 | Private donors |
| 14 | Ramanna Fellowship, DST, India | 49 | Richard Winter Foundation |
| | **International grants** | | **International govt. funding** |
| 15 | Alexander von Humboldt Foundation | 50 | Coastal Zone Management Centre, The Netherlands |
| 16 | American Society of Primatologists | 51 | French Academy of Agriculture |
| 17 | BBC Wildlife Fund | 52 | German Academic Exchange Service |
| 18 | Darwin Initiative, UK | 53 | German Research Council |
| 19 | Earthwatch UK | 54 | Grafog Land Niedersachsen |
| 20 | Idea Wild | 55 | Ministry of Economy and Competitiveness, Spain |
| 21 | International Association for Bear Research and Management | 56 | Natural Sciences and Engineering Research Council of Canada |
| 22 | IUCN Save our Species | 57 | NSF, USA |
| 23 | Marine Conservation Action Fund | 58 | Save the Tiger Fund, National Fish and Wildlife Foundation |
| 24 | National Geographic | 59 | Spanish National Research Council |
| 25 | National Geographic Society Emerging Explorers | 60 | United States Fish and Wildlife Service |
| 26 | Ocean Park Conservation Fund, Hong Kong | 61 | United States Geological Survey |
| 27 | People Trust for Endangered Species | | |
| 28 | Pew Marine Fellowship Programme | | **Awards** |
| 29 | Rufford Small Grants Foundation | 62 | Oriental Bird Club Forktail Leica Award |
| 30 | Snow Leopard Trust | 63 | Oriental Bird Club WildWings Conservation Awareness Award |
| 31 | The Peregrine Fund | 64 | Whitley Fund for Nature |
| 32 | Wildlife Conservation Society, USA | 65 | Wings World Quest Women of Discovery |
| | **State govt. funding** | | **Others** |
| 33 | Forest Department of Tamil Nadu | 66 | Auckland Zoo |
| 34 | Government of Karnataka (Vision Group on Science & Technology) | 67 | Centre for Development and Finance, Chennai |
| 35 | Kerala State Council for Science, Technology and Environment, Govt. Of Kerala | 68 | Dorabji Tata Trust grants |
| 36 | West Bengal Forest Department | | |
| 37 | West Bengal State Zoo Authority | | |

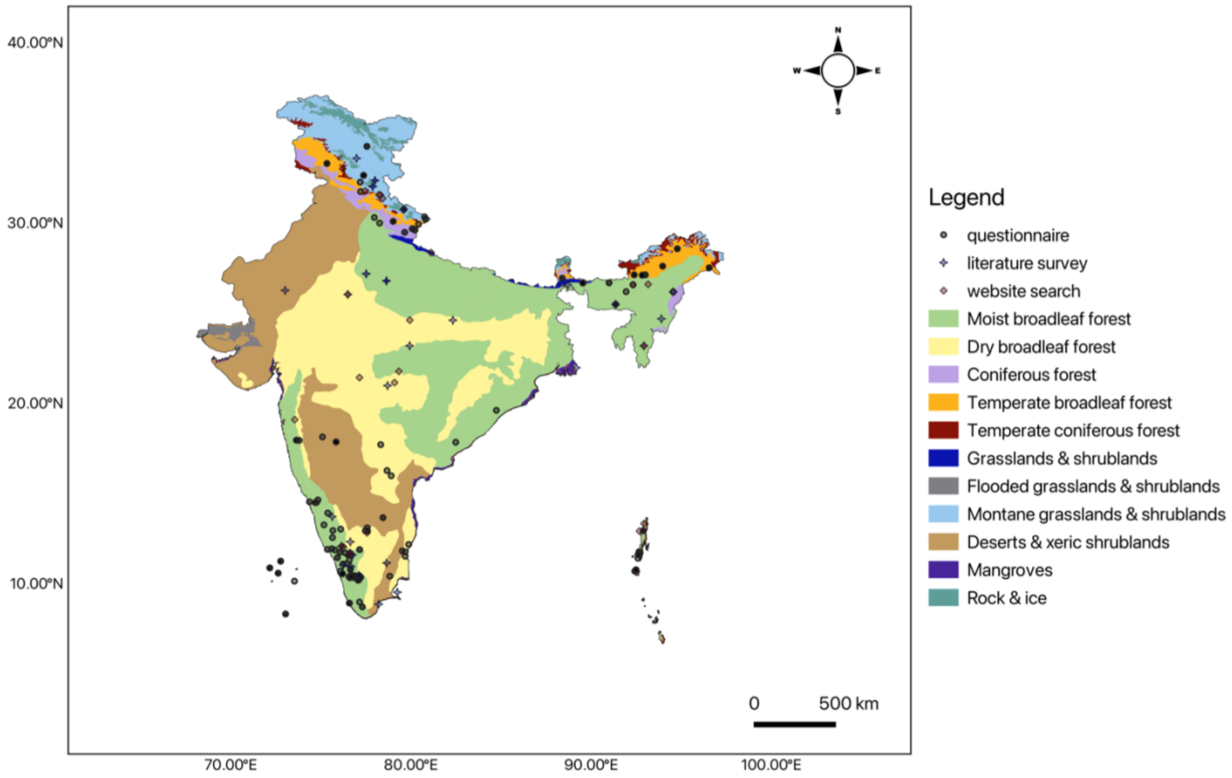

**Fig. SI1** Map of India showing the locations of LTEM efforts, with symbol shape and colour denoting source for LTEM site information. Grey circles: data from questionnaire survey, blue stars: literature survey, red diamonds: website search. Darker symbols are locations with multiple LTEM efforts, and overlapping symbols. Colours on the map are biomes.

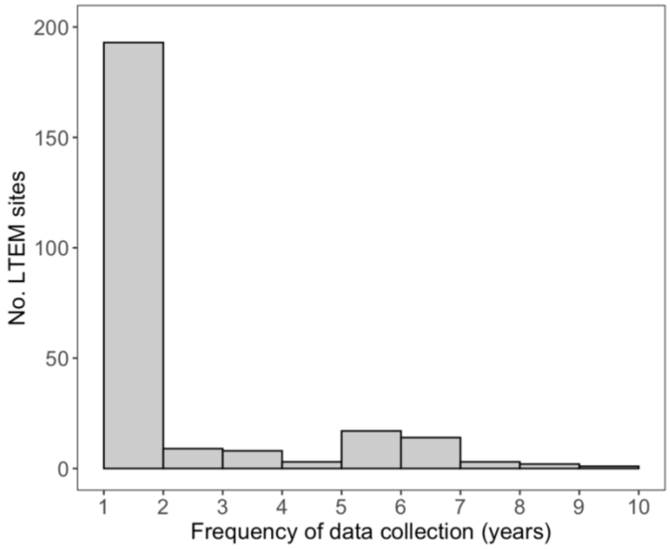

**Fig. SI2** Frequency of data collection in LTEM sites across India. Frequency of data collection was calculated as the number of years an LTEM effort has been ongoing divided by the number of years of data collection.

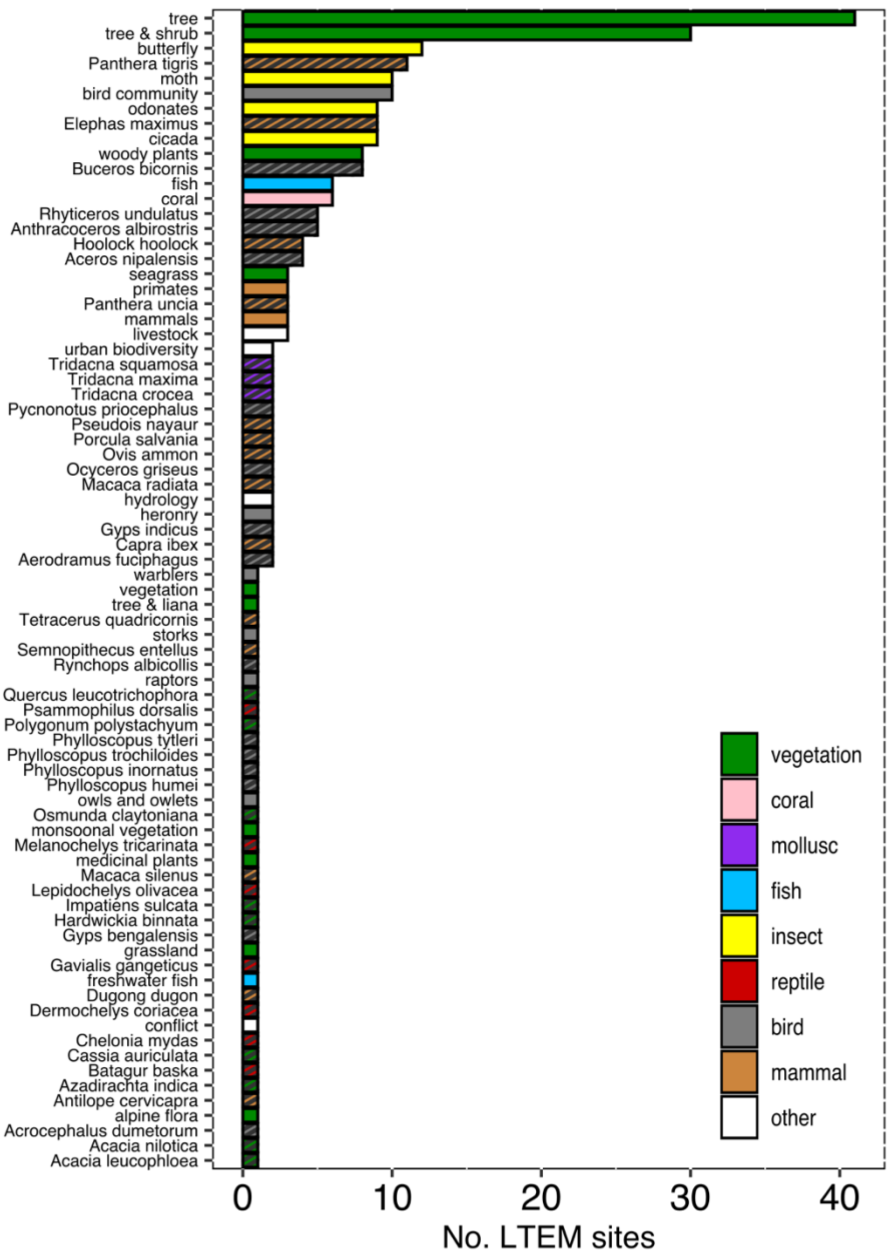

**Fig. SI3** Subjects of LTEM efforts in India. Colours are different ‘groups’ the subjects are classified under. Some LTEM subjects are species-specific (e.g. tiger under mammals); these are represented by striped bars. Some LTEM subjects are at the community level (e.g. primates under mammals); these are represented by plain filled bars. Biotic factors measured for each of these LTEM subjects may span ecological scales (Fig. 4). For instance, species-level biotic factors may be measured for community scale subjects (e.g. phenology data for dominant tree species in a forest monitoring LTEM). Community-level factors may be measured for species-specific LTEM subjects (e.g. prey species interactions in a tiger monitoring project).

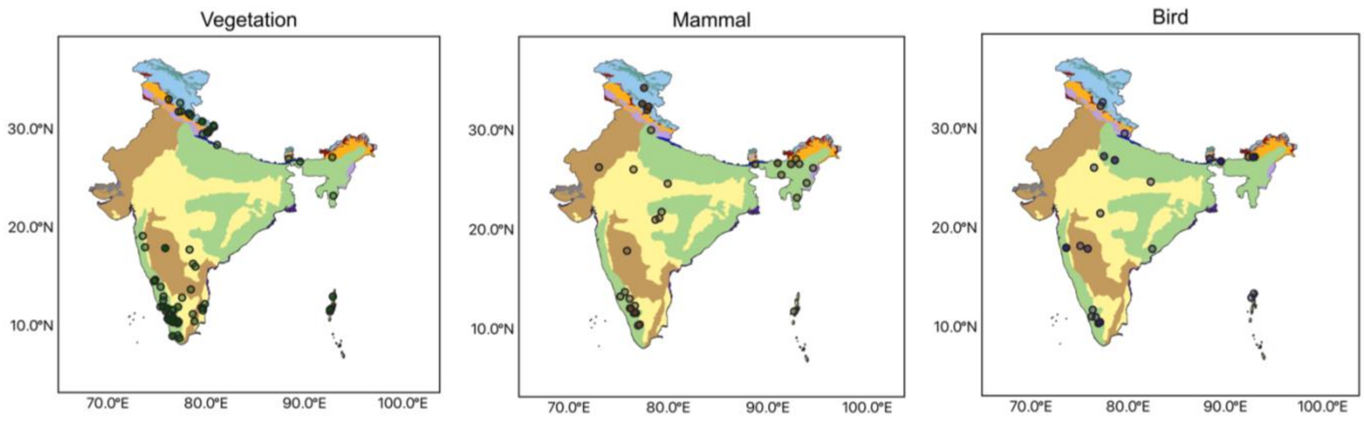

**Fig. SI4** Distribution of LTEM sites in India for some major groups of subjects monitored, to illustrate concentration of certain LTEM efforts in some regions more than others. Darker symbols are locations with multiple LTEM efforts, and overlapping symbols. Colours on the map are biomes (see legend in Fig. 1).

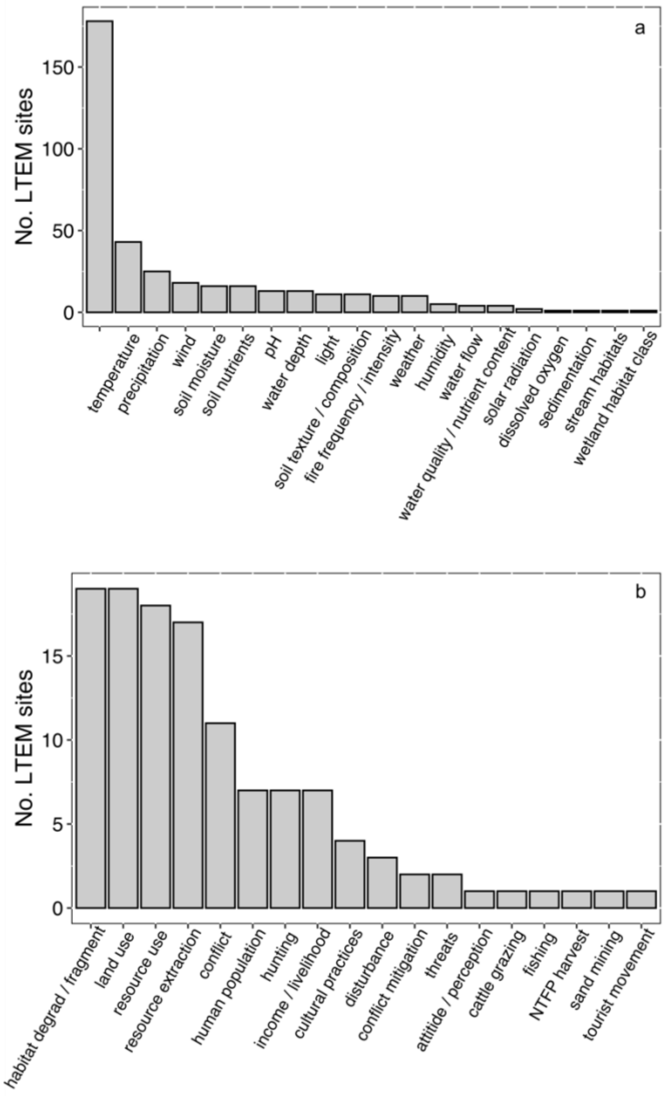

**Fig. SI5** (a) Abiotic and (b) Socioeconomic factors measured in the LTEM efforts.
